# Reduced glucose supply during neonatal infection attenuates neurological and renal pathology via modulation of innate and Th1 immunity

**DOI:** 10.1101/2024.05.15.594288

**Authors:** Jingren Zhong, Ole Bæk, Richard Doughty, Benjamin Meyer Jørgensen, Henrik Elvang Jensen, Thomas Thymann, Per Torp Sangild, Anders Brunse, Duc Ninh Nguyen

## Abstract

**Background:** Premature infants are highly susceptible to infections that can lead to sepsis with life-threatening organ dysfunctions. The clinical practice of high parenteral glucose supply in preterm infants can exacerbate infection outcomes through excessive glycolysis-induced inflammatory response. This in turn can affect the health of vital preterm organs, including the brain and kidneys. We hypothesized that reducing glucose supply in infected preterm newborns may help protect against pathology in these two key organs.

**Methods:** Caesarean-delivered preterm pigs were nourished with high or low parenteral glucose levels, infected with *Staphylococcus epidermidis* or saline, and cared for until 22h. Blood, brain, and kidney samples were collected at the end of the study for analyses.

**Results:** Infection led to multiple pathological changes, increased inflammation and tissue injury and dysfunction in both brain and kidneys of preterm piglets. Reduced glucose supply in infected animals alleviated neurological degradation, hyperemia and enhanced M2 microglial phenotype in the brain. This intervention also reduced plasma creatinine, renal edema, tubular vacuolization and dilatation. Multiple genes related to innate and Th1 immunity in both organs were highly correlated and dampened by reduced glucose supply, but there were clear signs that renal inflammation was closely connected to systemic inflammation while neuroinflammation was likely driven by immune response to the bacteria translocated into the brain.

**Conclusion:** Reduced glucose supply can protect brain and kidney health in infected preterm neonates.

## Introduction

Preterm birth (less than 37 weeks of gestation) accounts for 10% of all pregnancies [1]. Due to the compromised immunity and immaturity of multiple organ systems, preterm infants are particularly susceptible to neonatal infections (up to 40%) [2–4]. Neonatal infections can lead to sepsis, a state of organ dysfunction caused by an uncontrolled response to systemic infection that can have lifelong consequences, particularly neurodevelopmental sequelae [5, 6]. Upon clinical symptoms of neonatal infection, preterm infants are usually treated empirically with antibiotics [7]. However, the clinical signs of sepsis in preterm newborns are often subtle, and the utility of conventional biomarkers for infection is limited [8]. Likewise, prolonged use of antibiotics might be accompanied by side effects including disruption of normal immune development [9] and risks of toxicity to other organs, including the kidneys and central nervous system [10, 11]. Therefore, the development of supportive infection therapies could be crucial in preventing life-threatening sepsis in the context of reduced overall antibiotic use.

The neonatal brain and kidney are among the most susceptible organs to systemic inflammation [12, 13]. The blood-brain barrier is fragile in preterm infants and may be further disrupted during neonatal infection, resulting in an increased neuroinflammatory response that causes brain injury [14–16]. This neuropathology related to neonatal sepsis may lead to neurodegeneration and long-term consequences for cognition [17–19]. Likewise, the kidney is an organ with high perfusion rate and therefore highly susceptible to the consequences of systemic infection and inflammation [20, 21]. Sepsis is associated with altered hemodynamics, glomerular hypoperfusion, oliguria, dysregulated inflammatory responses, and oxidative stress-related tubular dysfunction in the kidneys [22–25]. In fact, sepsis is one of the leading causes of acute kidney injury (AKI) in clinically ill patients, including preterm infants [26, 27].

Due to intestinal immaturity, preterm infants often do not tolerate full enteral feeding, necessitating the use of parenteral nutrition (PN). To minimize the risk of hypoglycemia and hypoglycemia-induced brain injury, preterm infants often receive high parenteral glucose supply (14-17 g/kg/d) [28]. However, high parenteral glucose intake is linked to hyperglycemia, a condition that is associated with prolonged ventilator dependence and prolonged hospital stay in preterm septic neonates [29, 30]. In addition, there are no detailed guidelines for parenteral glucose supply in infants with suspected infection.

The negative effects of a high glucose intake during infection may be related to exaggerated glycolysis in immune cells, providing them with excessive fuel for inflammatory responses while disabling cell restorative functions [29, 31–34]. Using a neonatal sepsis model in *S. epidermidis*-infected preterm pigs [35, 36], we have previously shown that high parenteral glucose intake increases the risk and severity of neonatal sepsis with uncontrolled systemic inflammation, partly via increasing immune cell glycolysis and lactic acidosis [37]. In another recent follow-up study, reducing the glucose supply to half of what preterm infants often receive (5%, 7.2 g/kg/day), not only prevented systemic hyper-inflammation and sepsis, but also maintained normoglycemia in preterm pigs [38]. Thus, reduced parenteral glucose delivery during infection may be beneficial in preterm infants. However, it is unknown how this might affect other vital organ functions in both uninfected and infected neonates. Especially considering that glucose supply has a major impact on the metabolism and development of several organs, including the brain and kidneys [39]. In the current study, we hypothesized that the sepsis-protective low glucose regime could also prevent neurological and renal pathology in infected preterm pigs. Using histologic assessments and gene expression analyses, we examined pathologic outcomes related to innate and adaptive immune responses in these two organs.

## Materials and method

### Animal experimental procedures

All animal experiments were approved by the Danish National Committee on Animal Experimentation (2020-15-0201-00520). An overview of the study is shown in **Figure 1A**. Fifty preterm pigs (Duroc x Yorkshire x Danish Landrace, at day 106 of gestation, ∼90%) from 3 litters were delivered by cesarean section in accordance with a previously established protocol [40]. Immediately after birth, the preterm newborn pigs were resuscitated and fitted with umbilical arterial catheters and then randomized into 4 groups in a two-by-two factorial design: with/without *Staphylococcus epidermidis* infection (SE/CON) and nourished with high/low levels of glucose in PN (HIGH/LOW, 21% - 30 g/kg/d or 5% - 7.2 g/kg/d, respectively). Hence, four experimental groups were set: CON-HIGH (n=8), CON-LOW (n=9), SE-HIGH (n=16), and SE-LOW (n=17). After catheterization and initiation of the different PN regimens, animals were inoculated with either live *S. epidermidis* (SE) bacteria (1×10^9^ CFU/kg) or sterile saline through the arterial catheter as described previously [16]. Animals were closely monitored for any signs of sepsis, pain, and circulatory collapse for 22 h post-infection. Pigs with symptoms of septic shock during the experiment were immediately euthanized according to humane endpoints, which were pre-defined as blood pH < 7.1 accompanied by poor perfusion and deep lethargy [37]. At the end of the experiment, all pigs were sedated with Zoletil mix (0.1ml/kg), and blood was collected via cardiac puncture for biochemical analysis, followed by euthanasia via an intracardial injection of pentobarbital. Brain and kidney tissues were quickly collected after euthanasia. The hippocampus from the left hemisphere was removed, snap frozen, and stored at −80 °C for biochemical and q-PCR analyses. The entire right hemisphere was fixed in 4% phosphate buffered paraformaldehyde for histology. Kidneys were dissected longitudinally, then one half was fixed in paraformaldehyde for histology, and the other half was snap frozen and stored at -80℃ for q-PCR analyses.

**Figure 1.**
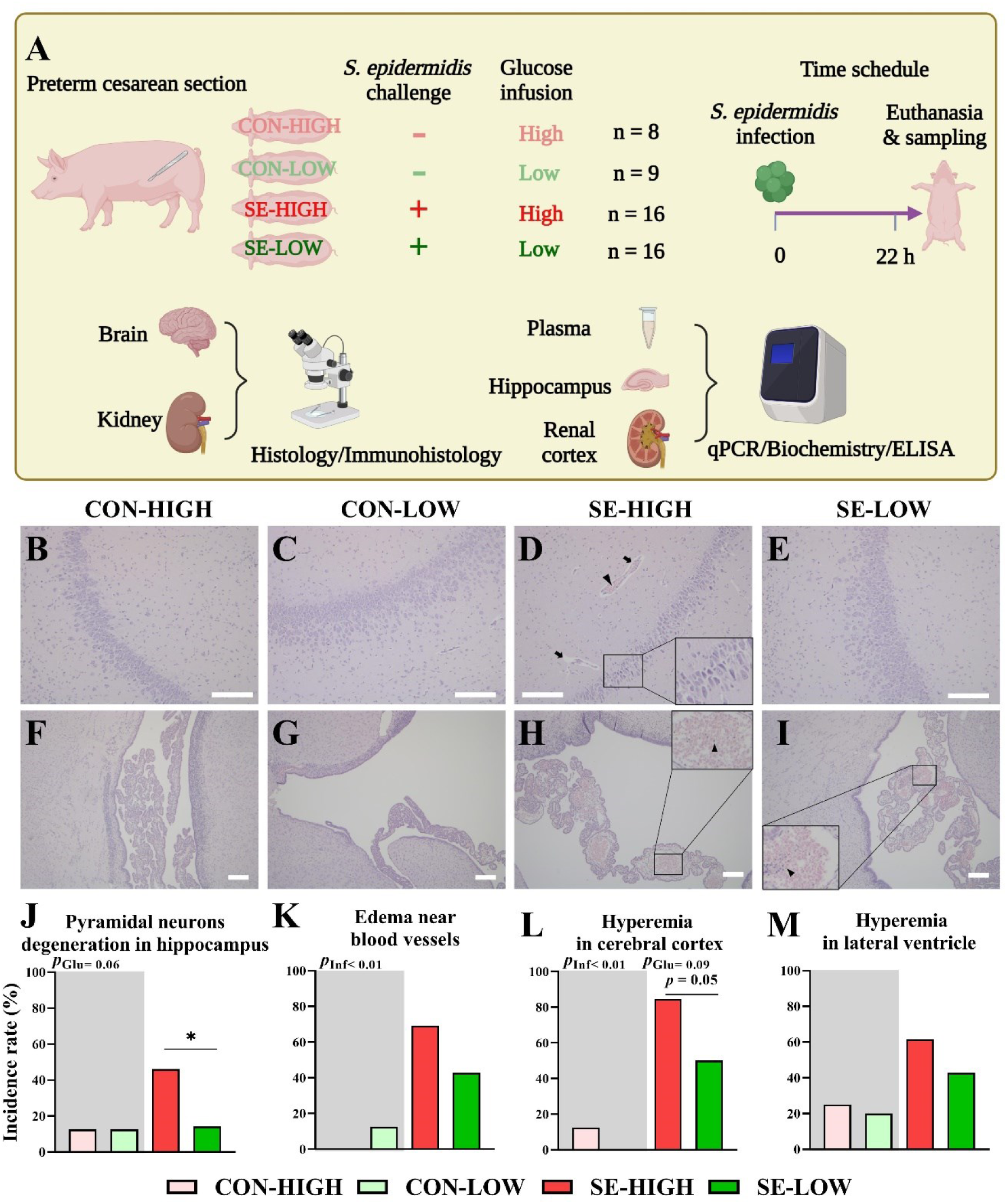
Study design and brain histopathology affected by infection and parenteral glucose supply. (A) Experimental design with newborn preterm piglets receiving PN with high (21%; 30 g/kg/d) or low (5%; 7.2 g/kg/d) glucose levels (HIGH/LOW), infected with 10^9^ CFU/kg S. *epidermidis* or saline (SE/CON) and cared for 22 hours before euthanasia or to the humane endpoints. Kidney and brain were analyzed for histology, qPCR and biochemistry. (B-I) Representative micrographs show histopathological symptoms in hippocampus (100 ×, B-E) and lateral ventricle (50 ×, F-I) (H&E staining, scale bars: 200 μm, n = 8-14 per group). Arrows indicate the edema happened near blood vessels. Arrowheads indicate hyperemia and infiltration of immune cells in the vascular lumen. The indicated square in Figure 1D shows the pyramidal neurons degradation in hippocampus. (J-N) The incidences of histopathological signs in the brain. SE, *S. epidermidis*. *P*_glu_ and *P*_inf_ indicate the significant impacts of glucose supply and infection, respectively, across all animals. CON, non-infected. SE, infected by *S. epidermidis*. HIGH/LOW, high/low parenteral glucose supply. *, **, *** *P* < 0.05, 0.01 and 0.001, respectively.

### Histological analysis

The fixed cerebral hemispheres were sectioned at a thickness of 5 mm (5 mm to 10 mm posterior to bregma) and embedded in paraffin [16]. The fixed kidney tissues were directly embedded in paraffin. All tissues were then sectioned at 4 μm, stained with hematoxylin and eosin (H&E), and evaluated for histopathological manifestations by blinded investigators. The neuropathological evaluation included the assessment of pyramidal neurons in the hippocampus, progenitor cell proliferation beneath the ependymal layer, neuron degeneration, edema in the cerebral cortex, hyperemia/vascular congestion in the lateral ventricle and cerebral cortex, and immune cell accumulation in vessels. The renal pathology evaluation included the examination of tubular dilatation, tubular vacuolization, interstitial edema, interstitial hemorrhages, and glomerular capillary hemorrhage. The histopathological signs were evaluated according to a 4-point scale (none; mild; moderate; and marked) or 2-point scale (none; present) based on the expertise of the two independent evaluators. Additionally, morphometry was performed on several kidney development-related parameters, including nephrogenic zone width (NZW), glomerular area, and density.

Brain sections were immunochemically stained for IBA-1, a marker of microglial cells [16, 41]. Following deparaffinization and rehydration, sections were treated with 1% hydrogen peroxide and the Biotin Blocking System X0590 (Dako, Glostrup, Denmark), blocked in 5% normal swine serum, and then incubated with 0.1% goat anti-human IBA-1 antibody. The sections were incubated with Ab5076 (overnight, 4 ℃; Abcam, Cambridge, United Kingdom) and treated with 0.05% biotinylated F(ab)’2 polyclonal rabbit anti-goat antibody (1 h, 20℃; E0466, Dako). Finally, the sections were treated with the Vectastain Elite ABC Kit (Vector Labs, Peterborough, United Kingdom) and 0.04% 3,30-diaminobenzidine (DAB; Sigma-Aldrich), and counterstained with hematoxylin. The digital images for the hippocampal region of the sections were captured under 100 × magnification using a slide scanner (Hamamatsu, NanozoomerS60, Hamamatsu, Japan). The images were analyzed using the ImageJ software (version 1.50i; National Institutes of Health, Bethesda, MD) to calculate the area percentage, density, and average size of IBA-1-positive cells.

### Biochemistry

Serum creatinine levels were analyzed using an Advia 2120i Hematology system (Siemens Healthcare, Ballerup, Denmark). The estimated glomerular filtration rate (eGFR) was calculated and indexed to body weight based on plasma creatinine [42]. β-hydroxybutyrate (BHB) and glucose levels in the hippocampus were measured via an enzymatic assay kit (MAK041, Sigma-Aldrich) and a glucose measurement kit (MAK263, Sigma-Aldrich), respectively.

### Gene expression analysis by RT-qPCR

Gene transcription levels in the hippocampus and renal cortex were analyzed using RT-qPCR. The primers utilized in the test are presented in Supplemental Table S1. Briefly, total RNA was extracted from homogenized tissues using the RNeasy Mini Kit (Qiagen, Copenhagen, Denmark). The RNA concentration and purity were determined using the NanoDrop 2000 (Thermo Scientific, MA, USA). Subsequently, the RNA was converted into cDNA with the High-Capacity cDNA Reverse Transcription Kit (Thermo Fisher Scientific, United States). RT-qPCR was then performed using the QuantiTect SYBR Green PCR Kit (Qiagen) on a LightCycler 480 (Roche, Hvidovre, Denmark), with the relative expression of target genes normalized to the housekeeping gene HPRT1 [43].

### Statistics

All statistical analyses were performed using R Studio 3.6.1 (R Studio, Boston, MA, United States). qPCR, biochemical, and histomorphometry data were fitted into a linear mixed-effect model using the lme4 and multcomp packages. The fixed factors included in the model were glucose level, infection status, their interaction (glucose × infection), sex, and birth weight. The random factor was litter. Binary data were analyzed with a logistic regression model, with group/infection status/glucose level, sex and litter as the fixed factors. If the data did not fit the model well, binary data were subjected to Fisher’s exact test. Subsequent to the initial analysis, a post hoc comparison was conducted between groups within each infection status. This was achieved through the application of a pairwise t-test to the data, which had been fitted using similar models. Categorical score data were analyzed by the Kruskal-Wallis test between groups under each infection status. A *p*-value < 0.05 was regarded as statistically significant. Except for categorical data, the data were presented as mean ± SEM.

## Results

We have previously demonstrated that the reduced glucose supply decreased the severity of sepsis during neonatal infection with corresponding marked shifts in hepatic metabolism [38]. Here we attempted to characterize whether these systemic effects are observed in two of the most vital organs in preterm neonates, the brain and kidney.

### Reduced glucose supply during neonatal infection attenuates neuropathology

The histological examination of brain H&E slices revealed infected animals had significant increases in the incidences of edema near blood vessels, hyperemia, and immune cell accumulation in vessels, compared to the non-infected controls (**Figure 1K and L, Supplemental Figure S1B**). The two non-infected groups (CON-HIGH and CON-LOW) had similarly low incidences of signs of pathology. Between the two infected groups (SE-HIGH and SE-LOW), animals with reduced glucose supply showed lower incidences of pyramidal neuron degeneration in the hippocampus (*p* < 0.05), hyperemia in the cerebral cortex (*p* = 0.05), and a tendency to lower incidences of edema near blood vessels and hyperemia in the lateral ventricle (**Figure 1J-M**). No discernible differences were observed among the four groups with respect to the incidence of neuronal degeneration near vessels in the cerebral cortex and progenitor cell proliferation under the ependymal layer (**Supplemental Figure S1 A and C**).

### Reduced glucose supply reduces neuroinflammation and modulates microglial morphology

Previous studies have demonstrated that *S. epidermidis*-infected preterm pigs exhibit clear cellular and molecular immune responses in the hippocampus [16]. In the present study, we examined these targets and the impact of reduced glucose supply. Neonatal infection led to a significant elevation of all tested hippocampal genes related to chemokines and innate immunity (from 2 up to 10,000-fold, all *p* < 0.001, **Figure 2A-C**). A comparison of the two infected groups revealed that reduced glucose supply significantly downregulated the expression of *MMP8*, *MMP9*, *ICAM1*, *IL4R*, *TAGLN2*, *IL8*, *SAA*, *CXCL10* and *CXCL11* (*p* < 0.05). Additionally, a comparison of the non-infected animals revealed that CON-LOW had increased S100A8 levels compared to CON-HIGH (*p* < 0.05).

**Figure 2.**
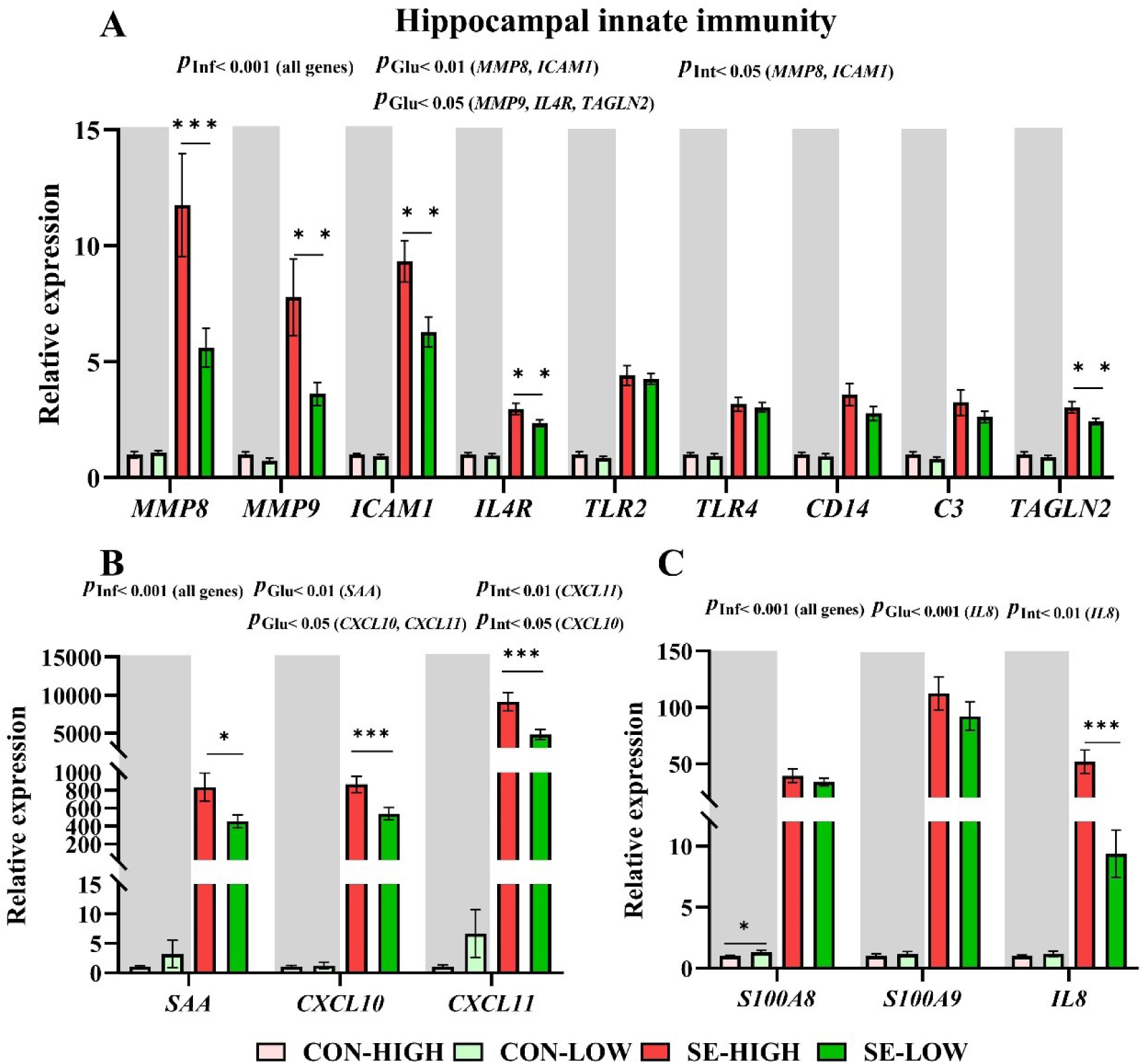
Effects of infection and glucose supply on hippocampal innate immune response. (A-C) Relative expression of genes related to innate immunity (n = 7-14/group). *P*_glu_ and *P*_inf_ indicate the significant impacts of glucose supply and infection, respectively, across all animals. *P*_int_ indicates significant interaction between glucose supply and infection. CON, non-infected. SE, infected by *S. epidermidis*. HIGH/LOW, high/low parenteral glucose supply. *, **, *** *P* < 0.05, 0.01 and 0.001, respectively.

We sought to determine whether the attenuation of neuroinflammation by reduced glucose supply was mediated via the activity of microglial cells. Hippocampal IBA1^+^ microglial cell staining (**Figure 3A-G**) clearly showed that infected animals had increased IBA1^+^ cell area percentage, density and average size (*p* < 0.001), indicative of an activated status with increased infiltration and enlarged cell bodies. This is in contrast to the resting status with small cell bodies and long and ramified processes in non-infected control animals [44]. Interestingly, SE-LOW animals exhibited higher levels of these microglial parameters than SE-HIGH animals (*p* < 0.01). Specifically, microglia in SE-HIGH animals had a rounded shape with relatively few processes while SE-LOW animals demonstrated a bushy shape with enlarged cell bodies and truncated processes. Given the decreased neuroinflammatory responses observed in SE-LOW vs. SE-HIGH, we further explored whether the distinct microglial morphology between these two groups was regulated by M1/M2 status. Most markers of M1/M2 microglia were upregulated by infection, with the exception of a downregulation observed for *TREM2* (*p* < 0.01). Furthermore, SE-LOW animals exhibited lower expression levels of both M1 (*IFNG*) and M2 (*IL6*, *ARG1*, *IL10*) targets than SE-HIGH animals (*p* < 0.05, **Figure 3H**). However, the ratio of the most well-known M1/M2 targets, *IFNG/IL10*, was lower in SE-LOW vs. SE-HIGH (*p* < 0.05, **Figure 3I**). Collectively, these data suggest that low glucose supply during neonatal infection may lead to relative more activation of M2 microglia, explaining less potent inflammatory responses and lower degree of pathologies in SE-LOW group. Furthermore, higher IL6 protein levels were expressed in the hippocampus of animals administrated with high glucose under both infected and normal conditions (*p* < 0.05, **Figure 3K**). However, no differences were observed in the level of IL10 (**Figure 3J**) and no TNF-α protein was detected in all the tissue (data not shown).

**Figure 3.**
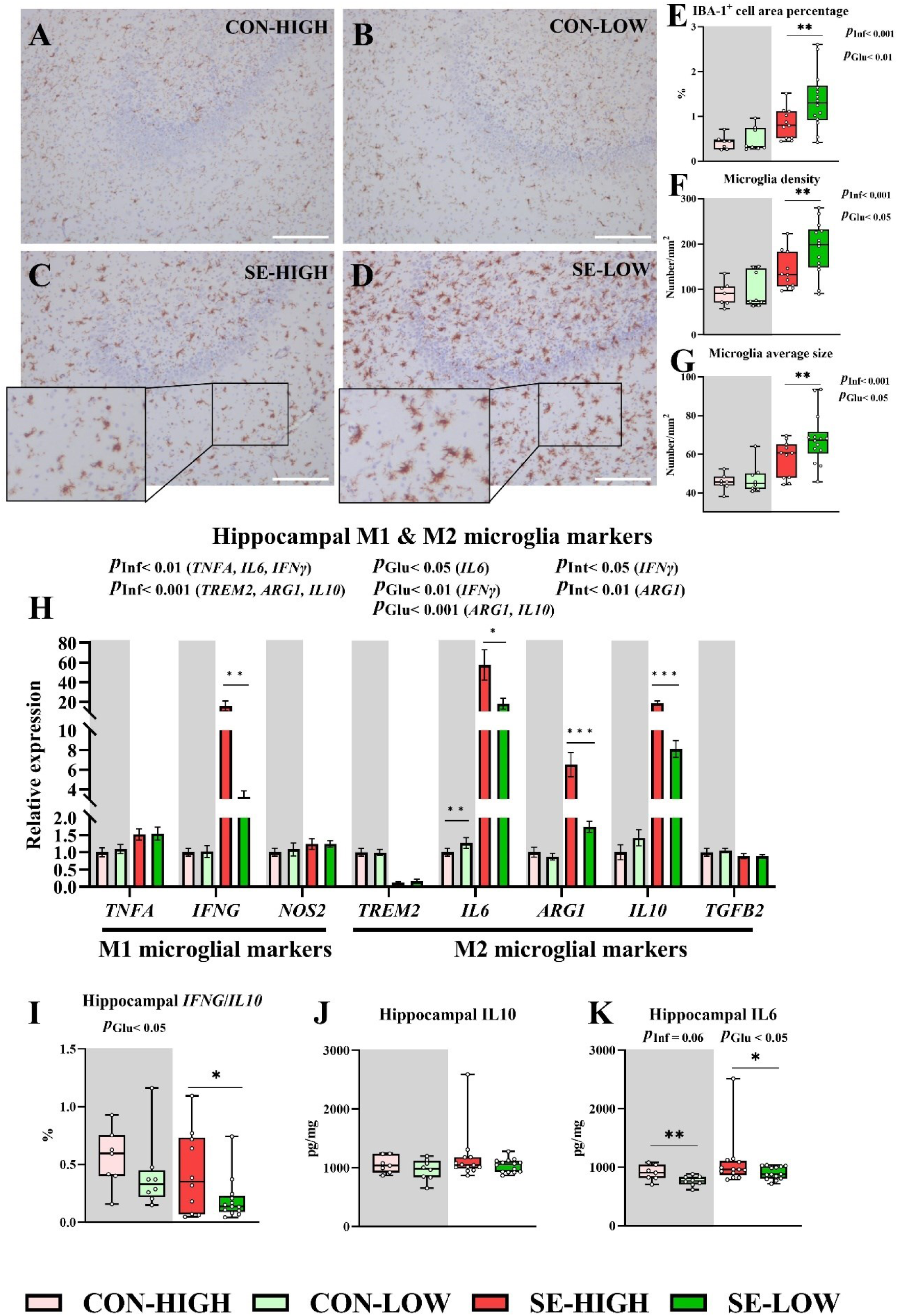
Hippocampal microglial activation and impacts of glucose supply and infection. (A-D) Representative micrographs show IBA-1^+^ stained microglial cells in the hippocampus (100 ×, counterstained with hematoxylin, scale bars: 200 μm). Microglia in two control groups (A and B) are characterized by small cell bodies with long and ramified processes. Microglia in SE-HIGH (C) show relatively rounded shape with few processes. Microglia in SE-LOW (D) show bushy shape with enlarged cell bodies and truncated processes. (E-G) IBA-1^+^ cell area (brown) percentage, density and average size (n = 7-14/group). (H) Relative expression of M1 and M2 microglia markers in the hippocampus (n = 7-14/group). (I) Ratio of *IFNG* to *IL10* as marker of M1/M2 ratio. (J and K) Hippocampal protein levels of IL10 and IL6, respectively. *P*_glu_ and *P*_inf_ indicate the significant impacts of glucose supply and infection, respectively, across all animals. *P*_int_ indicates significant interaction between glucose supply and infection. CON, non-infected. SE, infected by *S. epidermidis*. HIGH/LOW, high/low parenteral glucose supply. *, **, *** *P* < 0.05, 0.01 and 0.001, respectively.

### Glucose supply affects hippocampal metabolism and development

Neuropathology and inflammation are tightly connected with immune cell metabolism and neurodevelopmental status. In this study, we examined the expressions of targets related to these pathways in the hippocampus since this part of the brain is particularly susceptible to neurodegeneration in neuroinflammatory reactions and also responsible for critical functions like memory and vision [45, 46]. For metabolic targets, neonatal infection led to increased levels of *LDHA* (encoding the enzyme converting pyruvate into lactate during glycolysis) and *GLUT5* (encoding an exclusive glucose transporter mainly in microglia in the brain [47, 48], both *p* < 0.05, **Figure 4A**), suggesting that a high glucose supply increased glucose uptake into the hippocampal immune cells to fuel energy for inflammatory responses. Among the infected animals, those in the SE-LOW exhibited lower gene expression levels of glycolysis-related targets, including *HK1* (*p* < 0.05) and *LDHA* (*p* = 0.06), which is consistent with a reduction in inflammatory responses in these animals. Furthermore, both glucose and ß-hydroxybutyrate (*p* < 0.001, Figure 4B), a key ketone body and the alternative energy source to glucose for the brain, were higher in animals with a high vs. low glucose supply, irrespective of infection status (*p* < 0.06, Figure 4B and C).

**Figure 4.**
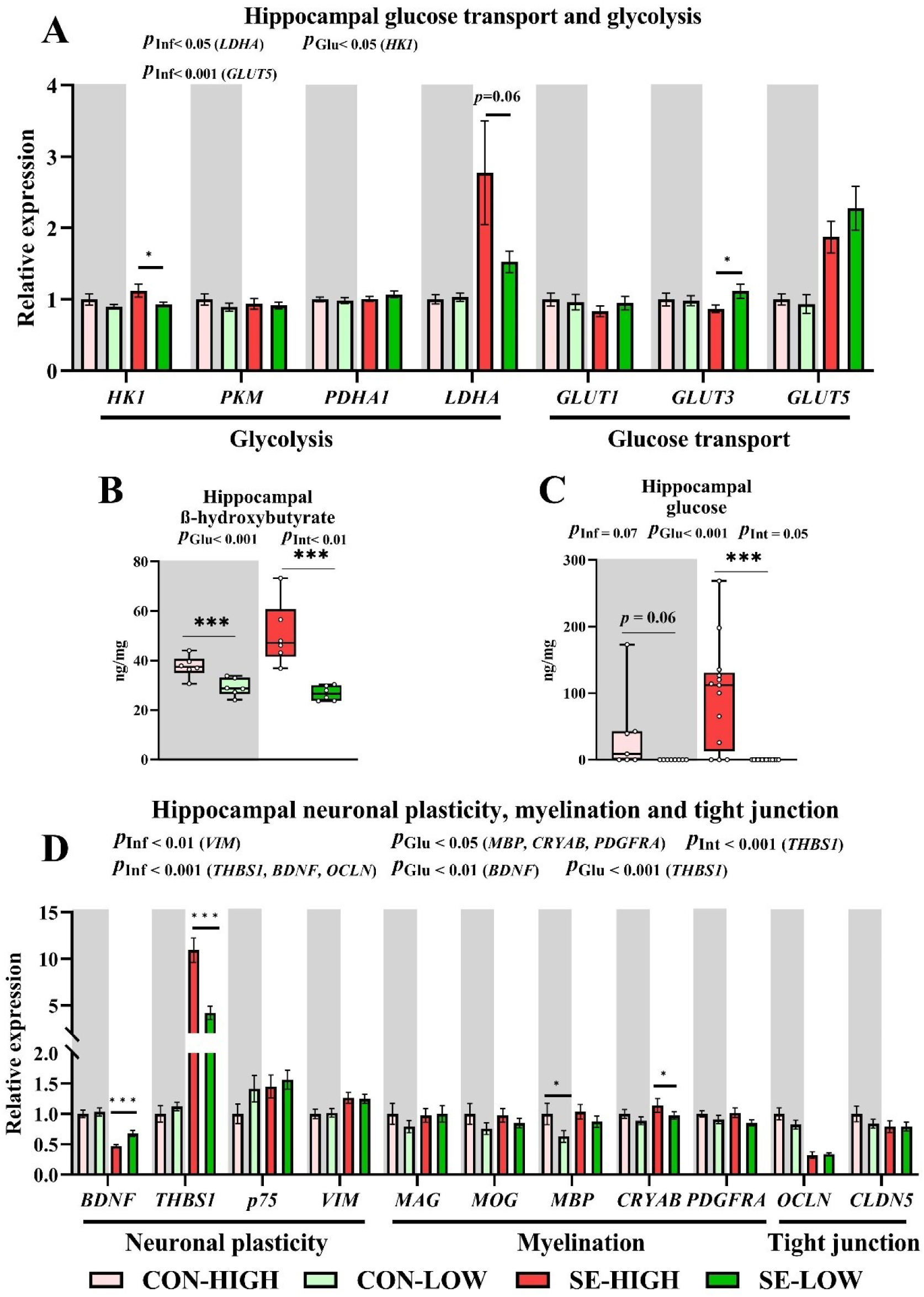
Effects of infection and glucose supply on hippocampal neuronal plasticity, myelination and metabolism. (A) Relative mRNA levels of genes related to glucose transport and glycolysis in the hippocampus (n = 7-14 per group). (B) ß-hydroxybutyrate concentration in the hippocampus (n = 6 per group). (C) Glucose concentration in the hippocampus (n = 6 per group); (D) Relative mRNA levels of genes related to neuronal plasticity, myelination and tight junction in the hippocampus (n = 7-14 per group). *P*_glu_ and *P*_inf_ indicate the significant impacts of glucose supply and infection, respectively, across all animals. *P*_int_ indicates significant interaction between glucose supply and infection. CON, non-infected. SE, infected by *S. epidermidis*. HIGH/LOW, high/low parenteral glucose supply. *, **, *** *P* < 0.05, 0.01 and 0.001, respectively.

As for neurodevelopmental targets, infection resulted in a downregulation of *BDNF* and an upregulation of *THBS1* and *VIM* (*p* < 0.01). Furthermore, reduced glucose supply during infection modulated *BDNF* and *THBS1* to the levels that were closer to the non-infected animals (*p* < 0.001, **Figure 4D**). The tight junction target *OCLN* was downregulated after infection (*p* < 0.001), without any impact of glucose supply. A few markers of myelination showed lower expression in animals receiving reduced glucose supply (*MBP*, *CRYAB*, *PDGFRA*, *p* < 0.05, **Figure 4D**).

### Systemic infection and glucose supply affect renal pathological outcomes

With the observed reduction in systemic and neuro-inflammation in infected animals with a reduced glucose supply relative to high glucose supply, we investigated how it affected the pathology in the kidneys as another infection-sensitive organ [20, 21]. Here, we observed a significant increase in serum creatinine and a decrease in eGFR in infected animals nourished with a high glucose supply compared to animals receiving a reduced glucose supply (*p* < 0.05, **Figure 5A and B**). This indicates a more pronounced degree of kidney dysfunction in SE-HIGH animals compared to SE-LOW animals. Subsequently, the extent of renal histopathological signs was evaluated. Following infection, more pronounced tubular dilatation and vacuolization, and higher incidence of interstitial edema were observed in SE-HIGH animals compared to SE-LOW animals (*p* < 0.01, **Figure 5C-E**). Minimal histopathological changes were observed in non-infected animals. The comparison of the incidences of commonly evaluated histopathological symptoms from the two independent evaluators is presented in Supplemental Figure S2. The incidence of the signs showed a similar trend between the two evaluators. The morphometric data of sections is shown in **Figure 5F-I**. Infection led to a significant decline in NZW, as well as in the area, density, and generation numbers of glomeruli (all *p* < 0.001). Conversely, a reduced glucose supply significantly restored all these structural parameters, bringing them closer to those observed in non-infected animals compared to a high glucose supply (*p* < 0.001).

**Figure 5.**
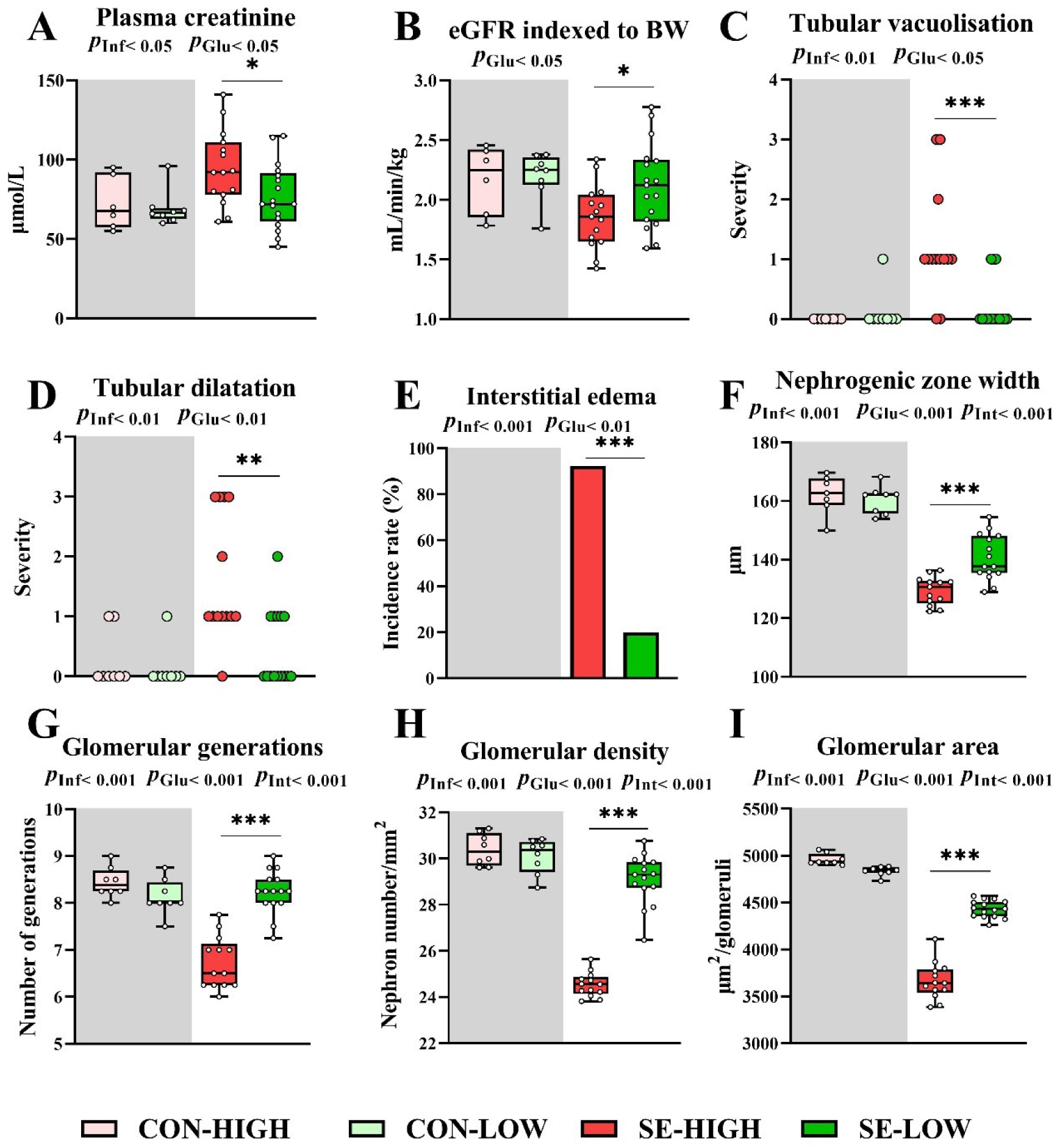
Effects of infection and glucose supply on kidney function, pathology and structure. (A) Plasma creatinine level; (B) Estimated glomerular filtration rate (eGFR) indexed to body weight (BW); (C and D) the pathological score of tubular vacuolisation and dilatation, respectively; (E) The incidence of interstitial edema; Morphometrical analysis of (F) the width of the nephrogenic zone; (G) medullary ray glomerular generation number; (H) glomerular density; (I) the cross-sectional area of glomeruli (n = 6-17 per group). *P*_glu_ and *P*_inf_ indicate the significant impacts of glucose supply and infection, respectively, across all animals. *P*_int_ indicates significant interaction between glucose supply and infection. CON, non-infected. SE, infected by *S. epidermidis*. HIGH/LOW, high/low parenteral glucose supply. *, **, *** *P* < 0.05, 0.01 and 0.001, respectively.

### Effects of glucose intake on renal inflammation and immunity

Next, we characterized the underlying cellular changes behind the pathological findings via gene expression analyses in the renal cortex, with targets related to innate and adaptive immune responses and cell metabolic pathways. The majority of the genes related to immunity were significantly upregulated by SE infection (*p* < 0.001), with the exception of *TNFA* and *GATA3* (**Figure 6A-B**). Relative to the SE-HIGH group, animals in SE-LOW showed downregulated expression of innate (*SAA*, *CXCL11*, *TLR2*, *TLR4*, and *C3*, *p* < 0.05) and adaptive immune genes (*IFNG*, *IL10* and *IL6, p* < 0.001). However, the levels of *LYZ* among the innate immune genes exhibited an opposite trend (*p* < 0.001). It is noteworthy that the key Th1 transcription factor *TBET* was not different between the two infected groups while the key Th2 transcription factor *GATA3* was up-regulated in the SE-LOW animals (*p* < 0.05). Th1/Th2 ratio targets were in general higher in infected vs. non-infected animals, but similar between two infected groups (**Figure 6C**). However, the Th1/Th2 ratios were both significantly higher in CON-LOW vs. CON-HIGH animals (*p* < 0.05). In line with the data related to the immune response, the expression levels of immune cell glycolysis markers (*PKM* and *HK1*) were also lower in SE-LOW vs. SE-HIGH animals (*p* < 0.05, **Figure 6D**). These data suggest that reduced glucose supply may have alleviated both innate and Th1 adaptive immune responses in the kidneys, possibly via the attenuation of immune cell glycolytic activity.

**Figure 6.**
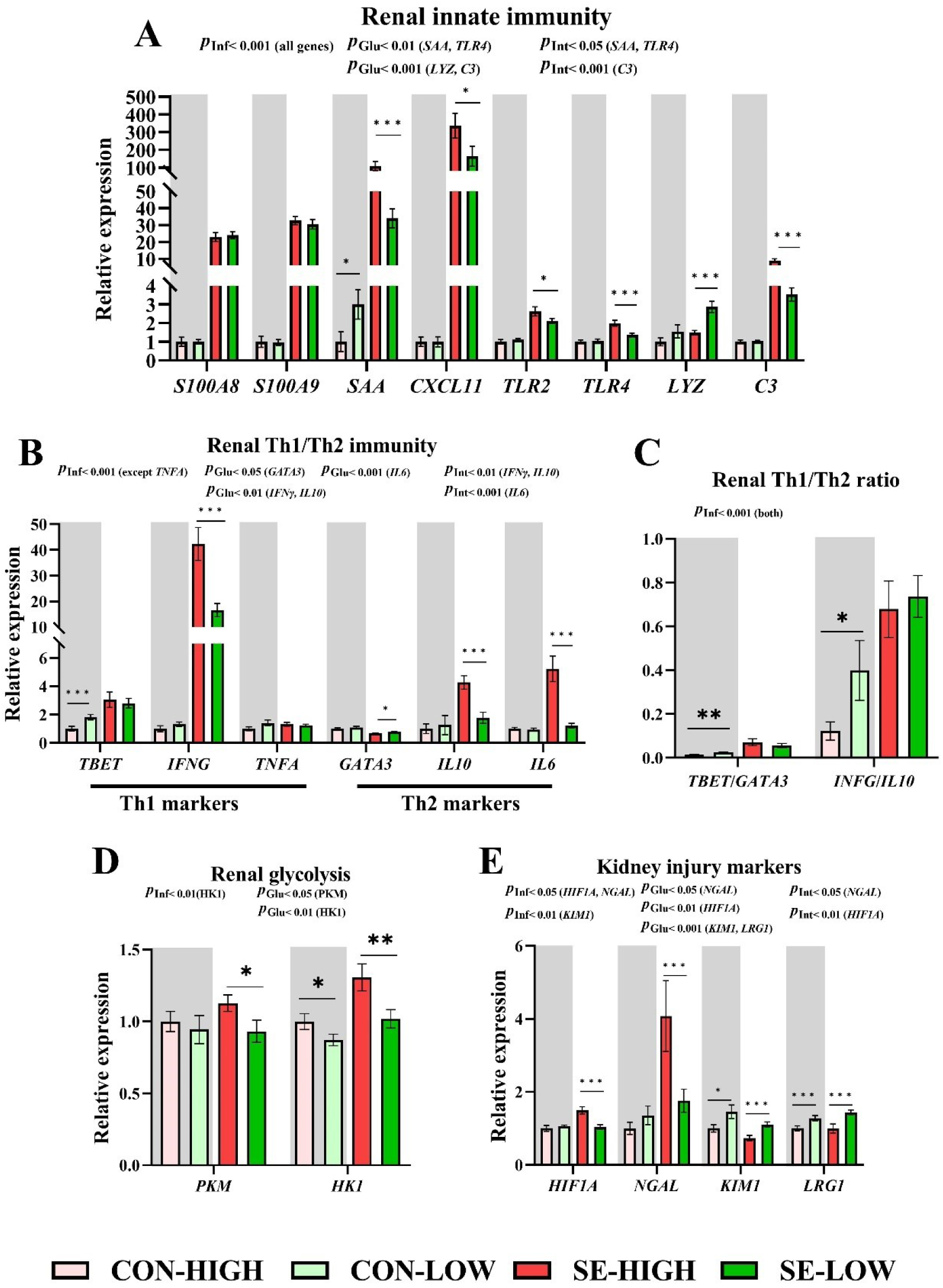
Effects of infection and glucose supply on renal cortex immune response. Relative expression of genes related to chemokines and innate immunity (A), Th1/Th2 immunity (B-C), glycolysis (D) and genes known as kidney injury markers (E) in the kidneys (n = 7-14 per group). *P*_glu_ and *P*_inf_ indicate the significant impacts of glucose supply and infection, respectively, across all animals. *P*_int_ indicates significant interaction between glucose supply and infection. CON, non-infected. SE, infected by *S. epidermidis*. HIGH/LOW, high/low parenteral glucose supply. *, **, *** *P* < 0.05, 0.01 and 0.001, respectively.

**Figure 7.**
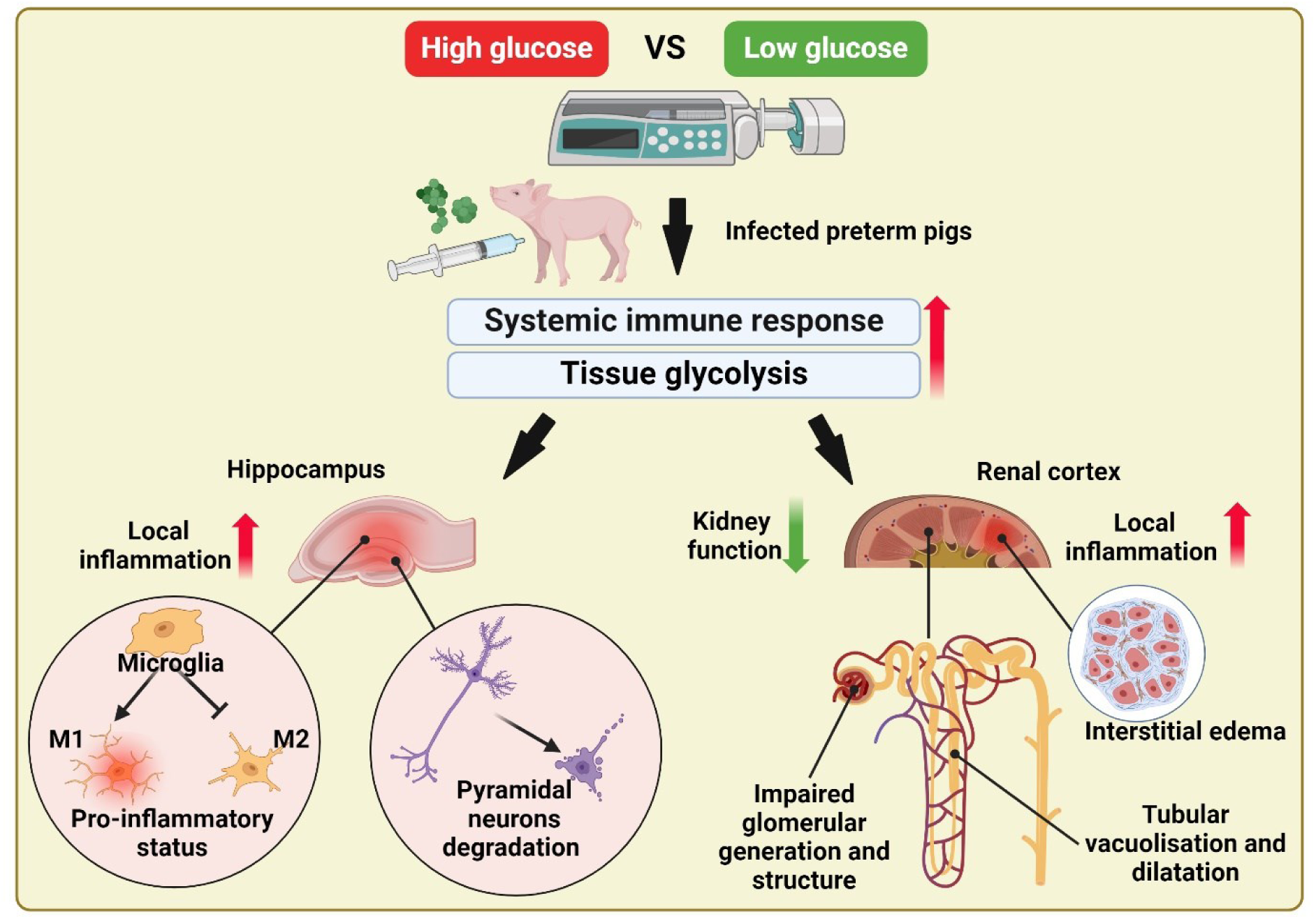
Summary of the study. In infected preterm pigs, compared to low glucose supply, high glucose supply induces stronger systemic inflammation and tissue glycolysis, which is further associated with increased local inflammation, M1 microglia polarization, and pyramidal neurons degradation in the hippocampus, and increased local inflammation, and impairment of renal structure and function.

We then further explored the gene expression of kidney injury markers (**Figure 6D**) and observed that SE-LOW animals exhibited lower expression levels of *HIF1A*, *NGAL* (*p* < 0.05), compared to the SE-HIGH animals, all in agreement with histological and immunity data. In contrast, two other kidney injury markers *KIM1* and *LRG1* demonstrated minimal changes in response to infection, as well as higher levels in animals with lower glucose supply, regardless of the infection status. This indicates that these markers are more influenced by glucose metabolism than injury in the context of neonatal infection.

### Correlation of the gene expression in brain and kidney, and plasma cytokines

To ascertain the potential connection of immune responses in the brain and kidney, a correlation analysis was performed on the gene expression between two organs (Table 1). The results indicated a positive correlation between the gene expression related to innate immune inflammatory responses in the kidneys (*S100A9*, *SAA*, and *TLR2*) and the brain (*CXCL10*, *CXCL11*, *ICAM1*, *S100A8*, *S100A9*, *TLR2*, *TLR4*, and *IL8*). Conversely, the expression of *S100A9*, *SAA*, and *TLR2* in the kidneys was negatively correlated with genes related to brain structure integrity (*OCLN*) and neural plasticity and development (*BDNF*). Next, we examined whether the gene expression of the two organs was correlated with systemic inflammation at euthanasia (Table 2). The overall plasma cytokine levels are shown in our previous study [38]. We found that the relative expression of most inflammatory genes and injury markers in the kidneys was positively associated with plasma cytokine levels. In comparison to the kidney, plasma cytokines exhibited a less pronounced positive correlation with the expression of inflammatory genes in the brain, with *IFNG* exhibiting no correlation. Therefore, kidney inflammation and pathology may be driven by the systemic inflammatory responses, whereas neuroinflammation and pathology may derive from a local immune response to the bacterial translocated into the brain.

**Table 1.**
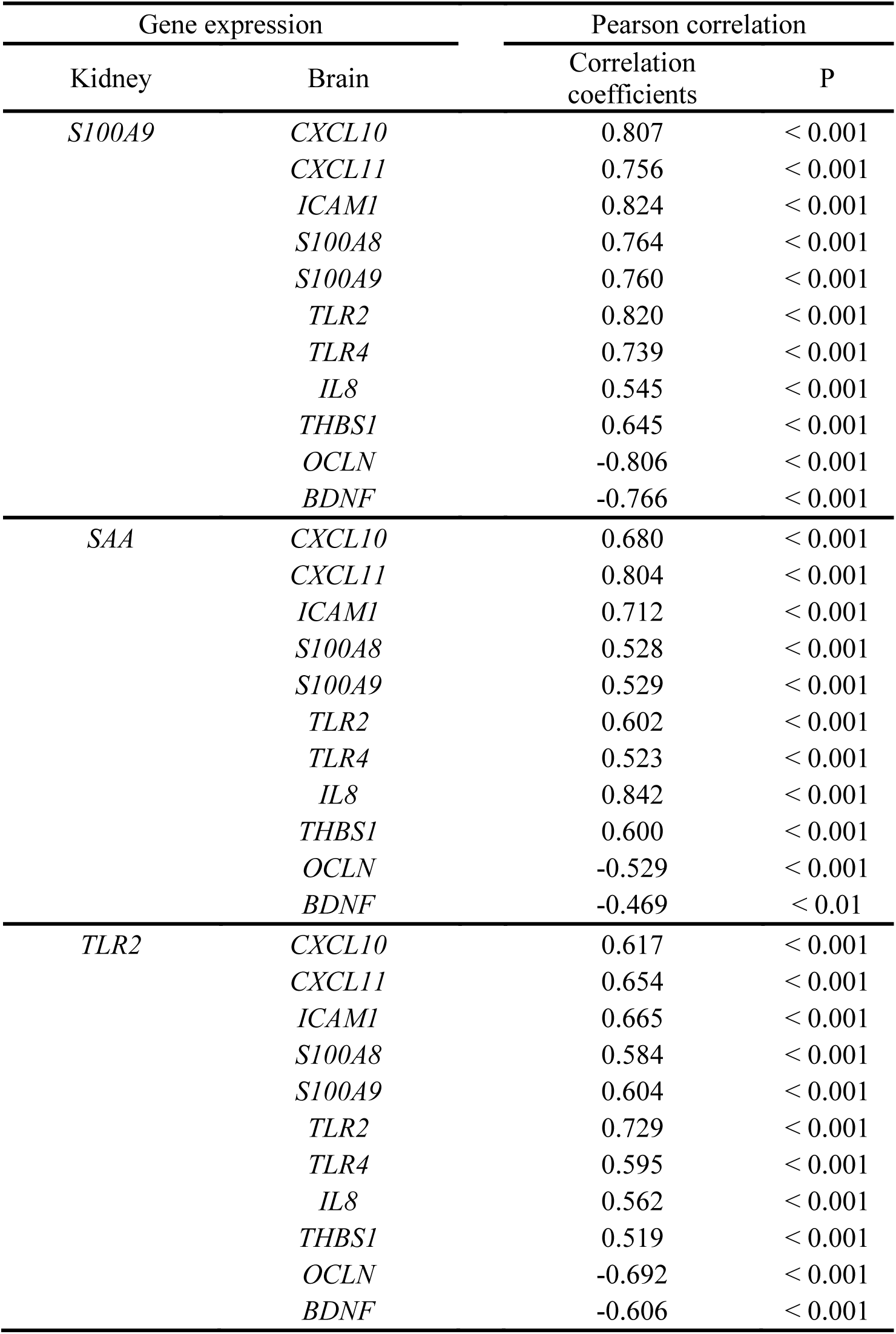
Pearson correlation of gene expression in kidney and brain.

**Table 2.**
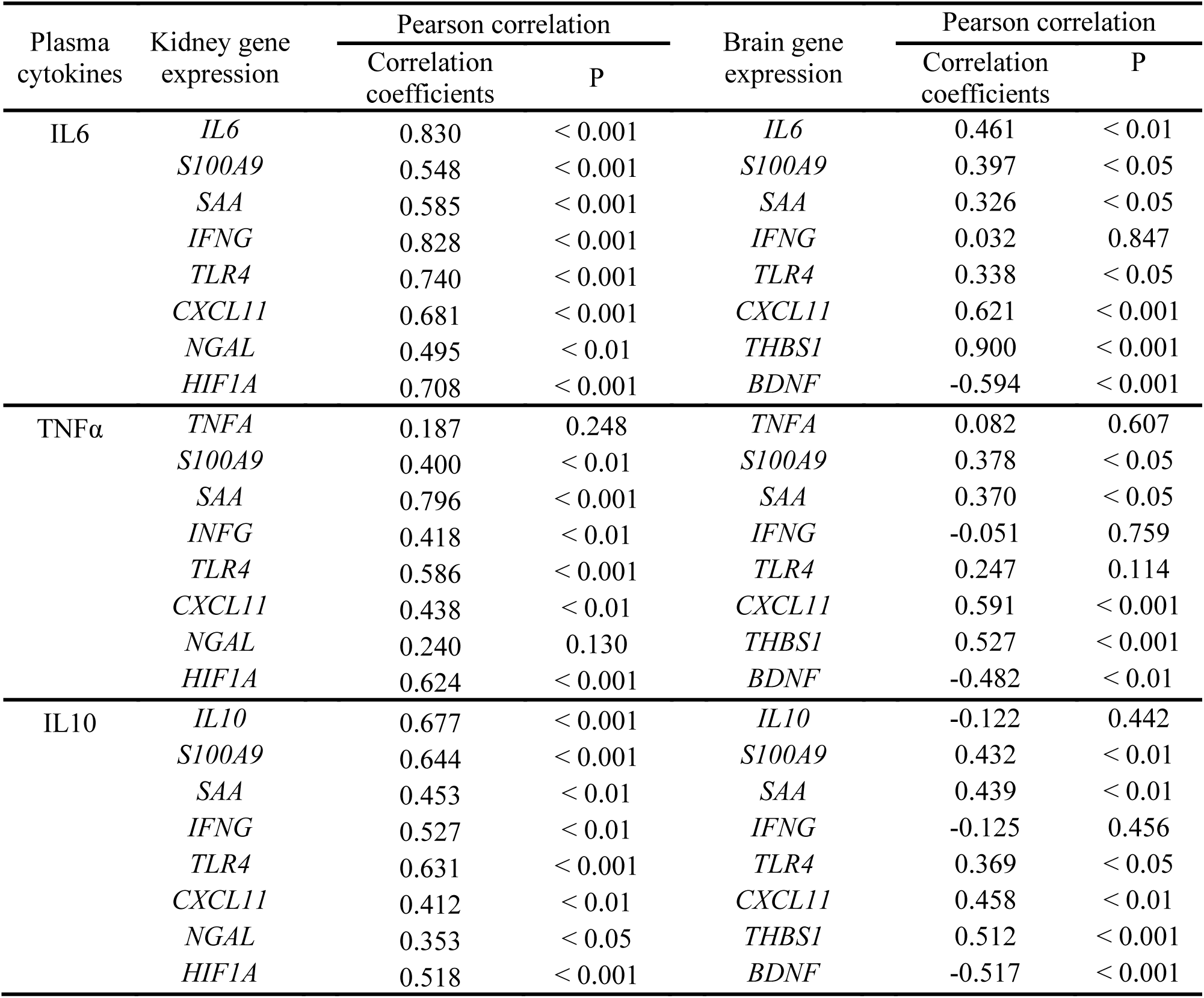
Pearson correlation of plasma cytokines and the gene expression in kidney and brain.

| Plasma cytokines | Kidney gene expression | Pearson correlation |  | Brain gene expression | Pearson correlation |  |
| --- | --- | --- | --- | --- | --- | --- |
|  |  | Correlation coefficients | P |  | Correlation coefficients | P |
| IL6 | *IL6* | 0.830 | < 0.001 | *IL6* | 0.461 | < 0.01 |
|  | *S100A9* | 0.548 | < 0.001 | *S100A9* | 0.397 | < 0.05 |
|  | *SAA* | 0.585 | < 0.001 | *SAA* | 0.326 | < 0.05 |
|  | *IFNG* | 0.828 | < 0.001 | *IFNG* | 0.032 | 0.847 |
|  | *TLR4* | 0.740 | < 0.001 | *TLR4* | 0.338 | < 0.05 |
|  | *CXCL11* | 0.681 | < 0.001 | *CXCL11* | 0.621 | < 0.001 |
|  | *NGAL* | 0.495 | < 0.01 | *THBS1* | 0.900 | < 0.001 |
|  | *HIF1A* | 0.708 | < 0.001 | *BDNF* | -0.594 | < 0.001 |
| TNFα | *TNFA* | 0.187 | 0.248 | *TNFA* | 0.082 | 0.607 |
|  | *S100A9* | 0.400 | < 0.01 | *S100A9* | 0.378 | < 0.05 |
|  | *SAA* | 0.796 | < 0.001 | *SAA* | 0.370 | < 0.05 |
|  | *IFNG* | 0.418 | < 0.01 | *IFNG* | -0.051 | 0.759 |
|  | *TLR4* | 0.586 | < 0.001 | *TLR4* | 0.247 | 0.114 |
|  | *CXCL11* | 0.438 | < 0.01 | *CXCL11* | 0.591 | < 0.001 |
|  | *NGAL* | 0.240 | 0.130 | *THBS1* | 0.527 | < 0.001 |
|  | *HIF1A* | 0.624 | < 0.001 | *BDNF* | -0.482 | < 0.01 |
| IL10 | *IL10* | 0.677 | < 0.001 | *IL10* | -0.122 | 0.442 |
|  | *S100A9* | 0.644 | < 0.001 | *S100A9* | 0.432 | < 0.01 |
|  | *SAA* | 0.453 | < 0.01 | *SAA* | 0.439 | < 0.01 |
|  | *IFNG* | 0.527 | < 0.01 | *IFNG* | -0.125 | 0.456 |
|  | *TLR4* | 0.631 | < 0.001 | *TLR4* | 0.369 | < 0.05 |
|  | *CXCL11* | 0.412 | < 0.01 | *CXCL11* | 0.458 | < 0.01 |
|  | *NGAL* | 0.353 | < 0.05 | *THBS1* | 0.512 | < 0.001 |
|  | *HIF1A* | 0.518 | < 0.001 | *BDNF* | -0.517 | < 0.001 |

## Discussion

The brain and kidneys are important organs in neonatal development and are especially susceptible to the effects of inflammation, particularly in preterm infants. The present study clearly demonstrated that SE infection in newborn preterm pigs resulted in evident signs of neurological and renal inflammation and related pathology. Besides, the study also revealed that a low parenteral glucose supply effectively alleviated the pathological findings, suggesting that approach could serve to reduce sequelae following serious early life infections.

Both the brain and kidney are susceptible to systemic infection and inflammation [49]. The brain is protected from blood-borne substances by the blood–brain barrier [50], while the kidney vasculature is highly fenestrated and thus more exposed to blood constituents [20, 21]. Examining the two organs not only allows us to monitor biological changes in key organs of preterm newborns under infection, but also increases our understanding of systemic infection impacts on organs of different types. In the present study, regardless of the blood–brain barrier, the brain of preterm pigs showed manifestations of local inflammation (cerebral cortex edema [51] and hyperemia [52]) and neuron impairment (pyramidal neurons pathological changes [53]) under neonatal bacterial infection. The histopathological changes demonstrate that neonatal systemic infection may lead to brain injury in preterm pigs, potentially associated with impaired blood–brain barrier function and increased levels of inflammatory mediators and bacterial products (as evidenced by the presence of IBA-1^+^ cells aggregates in the brain of infected animals, as shown in Supplemental Figure S1) [14]. The results were further supported by increased immune responses, compromised neuronal plasticity, and tight junction as indicated by gene expression data. In the kidney, neonatal infection resulted in glomerular filtration dysfunction, accompanied by renal tubular damage (tubular dilatation and vacuolization), parenchyma inflammation and interstitial edema. Increased innate and inflammatory responses, and more severe kidney injury were further supported by the gene expression data. It is noteworthy that despite the significant differences between the kidneys and the brain, the degrees of immune and inflammatory responses in the two organs exhibited a high correlation following infection. The immune reaction and health status of the two organs may be contingent upon systemic infection and immune responses. Nevertheless, further correlation analysis of systemic inflammatory markers with gene expression in the two organs indicates that the kidneys may still be more sensitive and susceptible to systemic responses, potentially due to its high perfusion rate.

When hosts are infected, aerobic glycolysis (as opposed to oxidative phosphorylation [31, 54]) becomes the main energy-generating process in immune cells for proinflammatory responses [31, 55, 56]. Neonates employ a tolerance defense strategy during serious infections, whereby they limit active immune responses [57, 58], partly by lowering glycolysis to avoid harmful hyperinflammatory and immunopathology [58, 59]. However, the clinical practice of sufficient and continuous glucose supply leads to hyperglycemia [30, 60], which might result in excessive immune cell glycolysis and disrupted pro- and anti-inflammatory balance [37]. In clinics, the negative effects of high glucose in preterm infants susceptible to infection include prolonged ventilator dependency, longer hospital stay, increased risk of sepsis and higher mortality rates [29, 33, 61]. **Supplemental Figure S3** presents an overview of gene expressions regulated by infection and glucose supply in the hippocampus and renal cortex. Notably, reduced glucose supply tends to attenuate the immune and inflammatory reactions induced by infection to a normal level in both the kidneys and hippocampus. Several glycolysis-related enzymes, which have been shown to promote inflammation in immune cells [62–65], were transcriptionally downregulated in the two organs in SE-LOW vs. SE-HIGH animals. The evidence above supports our hypothesis that the decreased inflammation and immune responses in animals receiving a reduced glucose level were related to a lower glycolytic process. However, it is important to note that the impacts of a high glucose supply may compromise the tolerance of preterm infants by multiple mechanisms other than excessive glycolysis. These include impairing neutrophil-mediated bacterial clearance [66], and facilitating biofilm formation, growth and adherence of the pathogen [67, 68]. Additionally, it is crucial to consider the importance of energy supply for normal neonate brain development. By measuring both the glucose and BHB levels in the hippocampus, it was observed that when glucose levels were decreased due to low glucose supply, the BHB, an important alternative fuel source, was also decreased in the brain. This decrease in BHB may be due to a more pronounced consumption of BHB as an alternative to glucose in the tissue. Thus, the longer-term consequences of a low glucose supply, after resolution of the inflammatory response, could be negative.

Newborn preterm neonates rely heavily on innate immunity for pathogen defense due to their limitations of the adaptive immune system [2, 69, 70]. However, the initial immunological milieu, skewed towards Th2 biased immune responses, are essential for newborn development [71]. In the brain, the polarization of microglia is key to maintaining neuronal health. The M2 phase of microglia facilitates repair and plasticity, while the M1 type activation of microglia can be deleterious [72]. Our gene expression data demonstrate a robust activation of inflammatory responses in both the brain and kidney following SE infection in pigs, while reduced glucose tended to dampen the inflammatory responses and concomitantly enhance protective M2 and Th2 pathways in the brain and kidney, respectively. The less severe injury conditions in SE-LOW vs. SE-HIGH animals were further corroborated by the relatively decreased expression of *THBS1* (produced in neuronal inflammation [73] and after brain injury [74]) and αB-crystallin (CRYAB) (exceedingly produced in brain lesions and involves in demyelination [75]) in the brain, and by the relatively decreased expression of kidney injury markers *HIF1A* and *NGAL* in the kidneys [76]. Notably, the protective effect of reduced glucose in the brain is also supported by the different morphology of activated microglia cells.

In SE-LOW animals, a greater number of microglia were in the intermediate activation status [44, 77], which attach to injured neurons for repair and can return to the normal resting state if no additional damage occurs [78]. In contrast, microglia in the SE-HIGH group were more in the phagocytic status (amoeboid) [44, 77]. The transformation into phagocytotic cells represents the final stage of M1 polarization [44, 79], which occurs under severe conditions such as cell death [78]. The evidence may also imply that reduced glucose supply potentially protects the brain from neuron dysfunction and severe injury associated with regulating microglia activation and polarization. However, it is currently unclear whether the intermediate status of macroglia is related to M2 activation. Interestingly in the kidney, *LYZ* was expressed higher in SE-LOW vs. SE-HIGH animals. The ability of lysozyme to kill internalized bacteria is important for phagocytic function. Our previous study demonstrated that SE-infection inhibited the lysosome pathway in pigs [37]. Therefore, it is possible that the reduced glucose intake assisted the host in defending against SE by enhancing the lysosome pathway, which is worthy of further investigation.

In non-infected animals, restricted glucose use led to a reduced ratio of Th2 to Th1 responses and increased *SAA* expression in the kidney, as well as a decrease in gene expression related to myelination process in the brain. These findings suggest that the level of glucose may affect neonatal physiology and the development of the two organs. While minimal histopathological changes were observed in non-infected animals supplied with restricted glucose, further detailed investigation is necessary to assess the impact of restricted glucose on normal organ development and physiology.

In summary, by using preterm pigs as our model, we demonstrated that a reduced parenteral glucose supply can benefit the brain and kidney health of preterm neonates under infectious conditions. This is achieved by modulating M1/Th1 immunity, alleviating innate immune responses, inflammation and tissue injury. However, under non-infectious conditions, sufficient glucose supply is still important for normal immune and tissue development. It is conceivable that a reduced parenteral glucose supply, during suspected infections in preterm neonates, could serve to protect them against organ dysfunction and to alleviate any possible sequalae.

## Acknowledgements

The authors highly appreciate experimental support from Jane Povlsen, Britta Karlsson and other group members from University of Copenhagen. The study was supported by local funding at the University of Copenhagen. The authors declare no potential conflicts of interest.

## Author contributions

DNN and PTS designed the study while OB contributed to the design. OB, AB and DNN performed animal experiments and tissue collection. JZ performed laboratory analyses and statistics. JZ, RD, BMJ, HEJ, TT, and AB contributed to histological/immunohistological examination. JZ, OB, AB and DNN contributed to data interpretation. JZ drafted the manuscript under supervision of AB and DNN. All authors contributed to manuscript revision and approved the final version.

