## Supplemental file for "Reduced glucose supply during neonatal infection attenuates neurological and renal pathology via modulation of innate and Th1 immunity"

**S Table 1 Primers for brain and kidney RT-PCR.**

| Gene | Gene full name | Forward | Reverse | Amplicon length |
|---|---|---|---|---|
| PPARA | Peroxisome Proliferator Activated Receptor Alpha | CCGAGACCGCAGATCTCAAG | GACGAAAGGCGGGTTATTGC | 128 |
| HK1 | Hexokinase-1 | TTTCCCTTGTCGGCAATCCA | CCTCCACTCCGCTTGCTTTA | 80 |
| PKM | Pyruvate Kinase M1/2 | GCCCTGGACACTAAAGGACC | CAGCCACAGGACATTCTCGT | 147 |
| LDHA | Lactate dehydrogenase A | GTCTGAAGGGTGGGGCATAC | AAATGCGACCAAAAGGGCAC | 157 |
| PDHA1 | Pyruvate dehydrogenase E1 subunit alpha 1 | GTCAGGAAGCTTGTTGCGTG | GGTAAAGCCATGAGCTCGGT | 86 |
| GLUT1 | Glucose transporter 1 (also SLC2A1) | ATCATCGGTGTGTACTGCGG | GTCCAGGCCAAATACCTGGG | 150 |
| GLUT3 | Glucose transporter 3 | GGAGCAAACAGGGTGTCACT | TGATCGCCTCAGGAGCATTG | 111 |
| GLUT5 | Glucose transporter 5 | GTGGTGCCCCAACTCTTCAT | TTGGCAAGGAGAGTCCGAAG | 74 |
| S100A8 | S100 Calcium Binding Protein A8 | AGGGAATTACCACGCCATCT | TTGAACCAGGTTTCTGCGTC | 99 |
| S100A9 | S100 Calcium Binding Protein A9 | GCCAAACTTTCTCAAGAAGCA | AGTGTCCAGGTCTTCCAGGAT | 70 |
| ICAM1 | Intercellular Adhesion Molecule 1 | CAATGTGGCCCCTAAACACCA | TGCATGGCAGAGTAGAGTGC | 194 |
| IL8 | Interleukin 8 (CXCL8) | CTGTGAGGCTGCAGTTCTGG | CCAGGCAGACCTCTTTTCCAT | 99 |
| MMP8 | Matrix Metallopeptidase 8 | ACTATGGCTTCCCGAGCAGT | AGGAAGAAGTGATGTTGCTGG | 205 |
| MMP9 | Matrix Metallopeptidase 9 | GGATGGGAAGTACTGGCGAC | ACACTTGGCGTCCAGAGAAG | 158 |
| SAA | Serum amyloid A | GCTAAAGTGATCAGCGATGC | AGTGGTTGGGGTCCTTGC | 145 |
| CXCL11 | C-X-C motif chemokine 11 | TTTGCATTGGCCCTGGAGTA | GCATCTTCGTCCTTTATGTGCT | 131 |

| | | | | |
|---|---|---|---|---|
| CXCL10 | C-X-C motif chemokine 10 | TCGCTGTACCTGCATCAAGA | GTGGGCAAGATTGACTTGCAG | 92 |
| TAGLN2 | Transgelin 2 | AGCGCTATGGCATCAACAC | ACACAGGCCATGTTCTTTCC | 73 |
| TLR4 | Toll Like Receptor 4 | TGGTGTCCCAGCACTTCATA | CAACTTCTGCAGGACGATGA | 116 |
| TLR2 | Toll Like Receptor 2 | CGTGTGCTATGACGCTTTCG | GTACTTGCACCACTCGCTCT | 232 |
| CD14 | Cluster of differentiation 14 | TCGAGGACCTGGAGGTAAC | CCCACGACACATTACGGAG | 96 |
| C3 | Complement C3 | ATCAAATCAGGCTCCGATG | GGGCTTCTCTGCATTTGATG | 76 |
| IL6 | Interleukin 6 | TGGGTTCAATCAGGAGACCT | CAGCCTCGACATTTCCCTTA | 116 |
| TNFA | Tumor Necrosis Factor alpha | ATTCAGGGATGTGTGGCCTG | CCAGATGTCCCAGGTTGCAT | 120 |
| iNOS/NOS2 | Nitric oxide synthase 2 | CAACAATGGCAACATCAGG | CATCAGGCATCTGGTAGC | 119 |
| IFN-γ | Interferon gamma | AGCTTTGCGTGACTTTGTGT | ATGCTCCTTTGAATGGCCTG | 247 |
| ARG1 | Arginase 1 | GTGGATGCTCACACCGACAT | GGGACCTCGGGAATCTTTTC | 113 |
| TREM2 | Triggering receptor expressed on myeloid cells 2 | CAATCTTCAAGCCCACGACG | GGGATTCAAGAGGGTCAGCC | 113 |
| TGFB2 | transforming growth factor beta 2 | GCGCGATTTGCAGACTTGAG | ATGTAAAGTGGACGCAGGCA | 170 |
| IL10 | Interleukin 10 | GTCCGACTCAACGAAGAAGG | GCCAGGAAGATCAGGCAATA | 73 |
| AIF1 | Allograft inflammatory factor 1 | ATACTCTGTCCCTGACCTGCC | AAGCCGAGAGAGGAAGCACT | 198 |
| MOG | Myelin Oligodendrocyte Glycoprotein | AGCGCAGACTGAGAGGAAAAC | CCGAGAACTGGCACGATCA | 121 |
| MAG | Myelin Associated Glycoprotein | CCGTGGGAAGAGAGTGTTGAA | AGTGAGCAACAGCTCCGTCT | 122 |
| MBP | Myelin Basic Protein | TGACTACAAACCGGCTCACA | TCCCAGCTTGAAGATTTTGG | 79 |
| CRYAB | Crystallin Alpha B | TCCCTGAGCCCCTTCTACTT | CATCTCTGAGAGCCCAGTGTC | 78 |
| BDNF | Brain Derived Neurotrophic Factor | CCCTACCCCTTCTTTTGACCA | TCTCACCTGGTGGAACTTTTC | 138 |

| | | | | |
|---|---|---|---|---|
| p75 | Nerve growth factor receptor (NGFR) | CTGCAAGCAGAACAAGCAAG | TCTGGCTGTCCACAGAGATG | 107 |
| THBS1 | Thrombospondin 1 | CAGAGAGACACCGACATGGA | GTCAAGCTGATCCGGATTGT | 78 |
| VIM | Vimentin | GTACCGGAGACAGGTGCAGT | CGTTCCAGAGACTCGTTGGT | 72 |
| PDGFRA | Platelet-derived growth factor receptor alpha | ACGCTACCAGTGAAGTCTACG | CGGGATGGTCACTCTTCAGG | 188 |
| OCLN | Occludin | CAGGTGCACCCTCCAGATTG | TGATTGGGTTTGAATTCATCAGGC | 146 |
| CLDN5 | Claudin 5 | CTGGACCACAACATCGTGAC | AGCACCGAGTCGTACACCTT | 107 |
| TBET | T-box transcription factor 21 | CTGAGAGTCGCGCTCAACAA | ACCCGGCCACAGTAAATGAC | 121 |
| GATA3 | GATA binding protein 3 | ACCCCTTATTAAGCCCAAGC | TCCAGAGAGTCGTCGTTGTG | 92 |
| LYZ | Lysozyme | TAAAGCATGGGTGGCATGGA | CAGTTTGCAACCCCGAATGT | 73 |
| HIF1A | Hypoxia Inducible Factor 1 Subunit Alpha | TGTGTTATCTGTCGCTTTGAGTC | TTTCGCTTTCTCTGAGCATTC | 96 |
| NGAL | Lipocalin-2 (LCN2) | AAGACGGCAGCTACAACGTC | GACACCACACGCACGACATA | 157 |
| KIM1 | hepatitis A virus cellular receptor 1(HAVCR1) | ATGTACCCTTGGGTAACCGC | AACGTAGAACATGCCCCTCG | 164 |
| LRG1 | Lipocalin-2 (LCN2) | AAGACGGCAGCTACAACGTC | GACACCACACGCACGACATA | 157 |

**Supplemental Figure S1. Pathological signs and IBA-1$^+$ stained microglial cells aggregates in the brain. (A-C)** The incidences of histopathological signs in the brain. (D) Representative micrographs show IBA-1$^+$ stained microglial cells aggregate in the brain (400 ×, counterstained with hematoxylin, scale bars: 50 μm). (E) The count of aggregates per slice. $P_{inf}$ indicates the significant impact of infection across all animals.

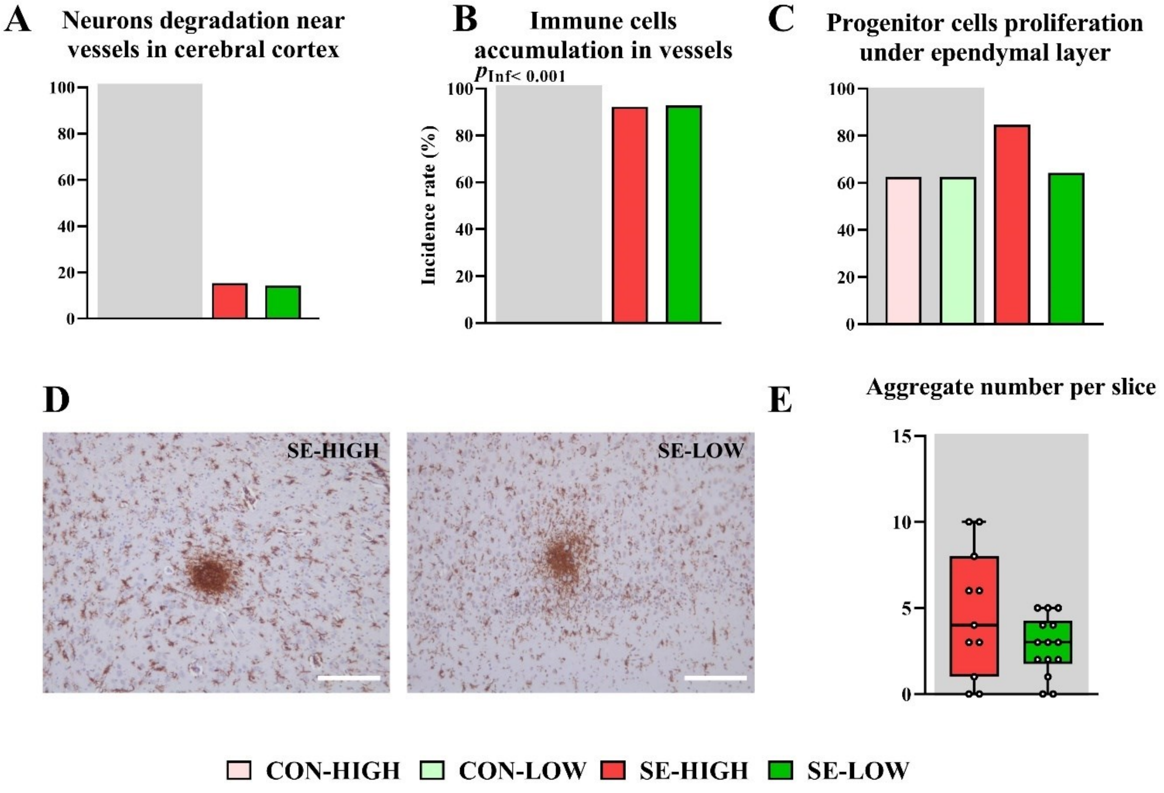

**Supplemental Figure S2.** Comparison of pathological evaluation of kidney H&E sections from two independent evaluators**.** The results are reported as the incidence of minimal pathological symptoms.

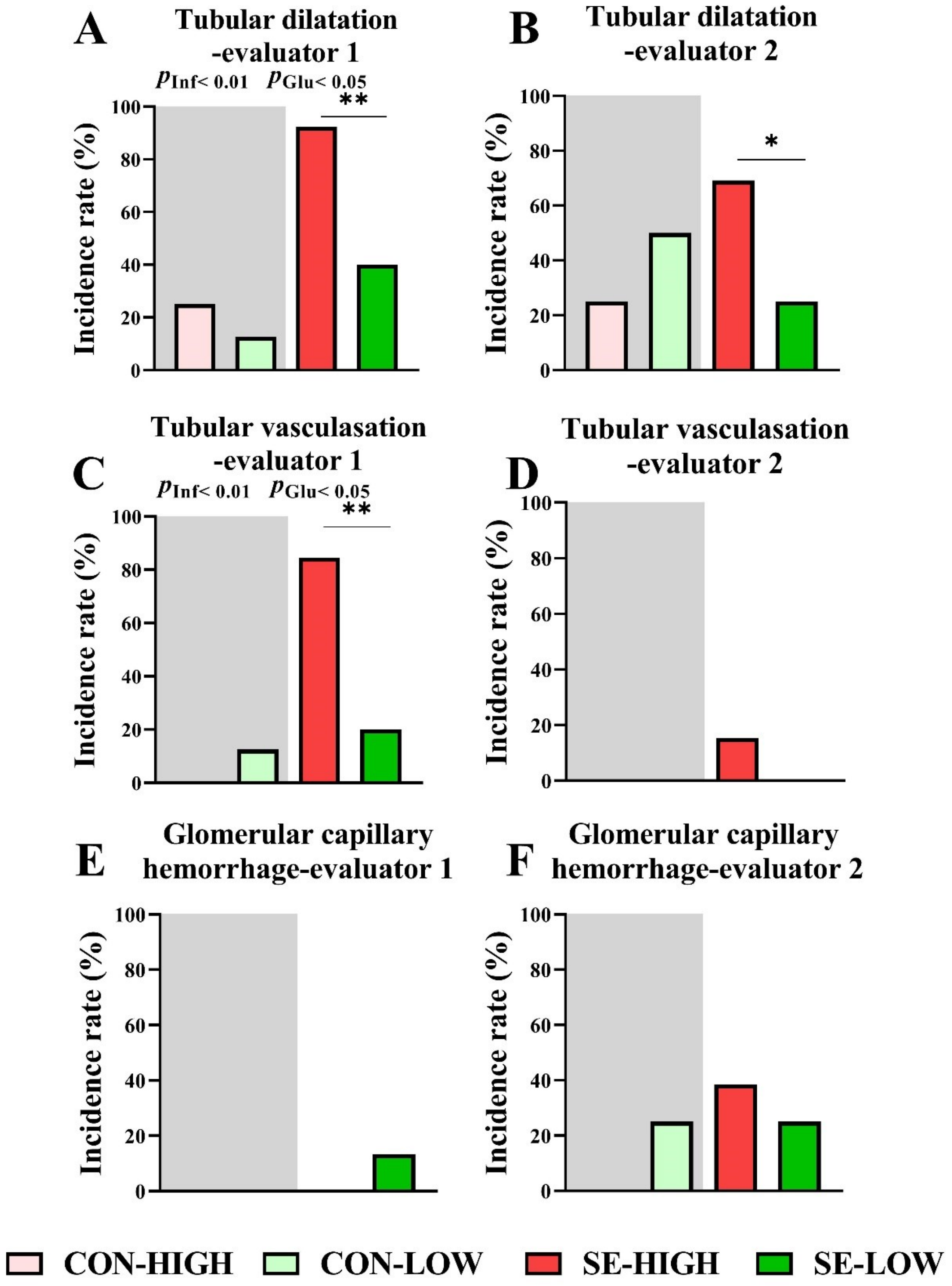

**Supplemental Figure S3. Overview of gene expressions regulated by infection and glucose supply in the hippocampus and renal cortex.** Gene expression in the hippocampus (A) and renal cortex (B). z-scores of expression level were depicted in colors from green (low) to red (high).

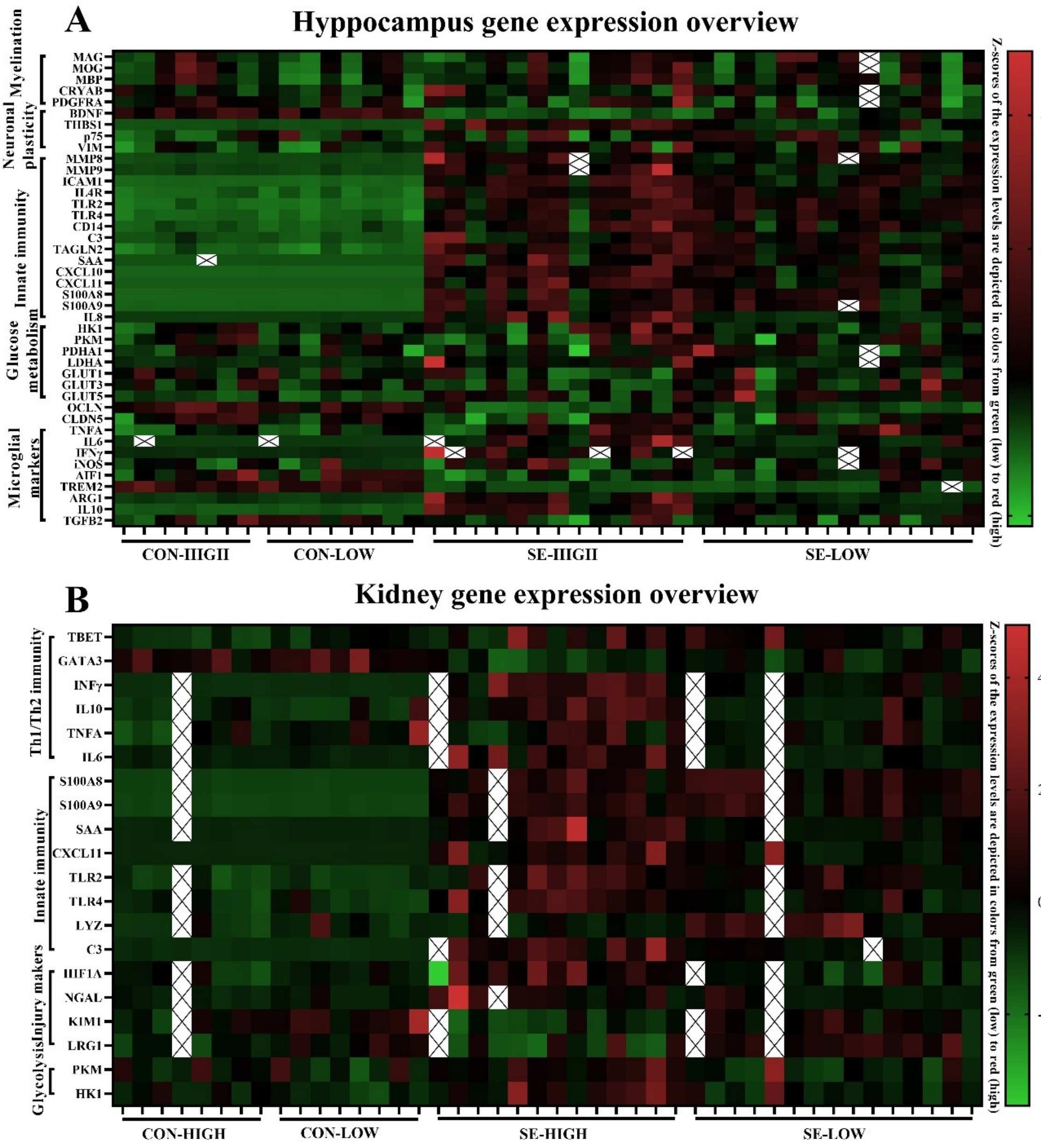


----
